## Supplementary material for "Direct binding of TDP-43 and Tau drives their co-condensation, but suppresses Tau fibril formation and seeding": Expanded View Figures

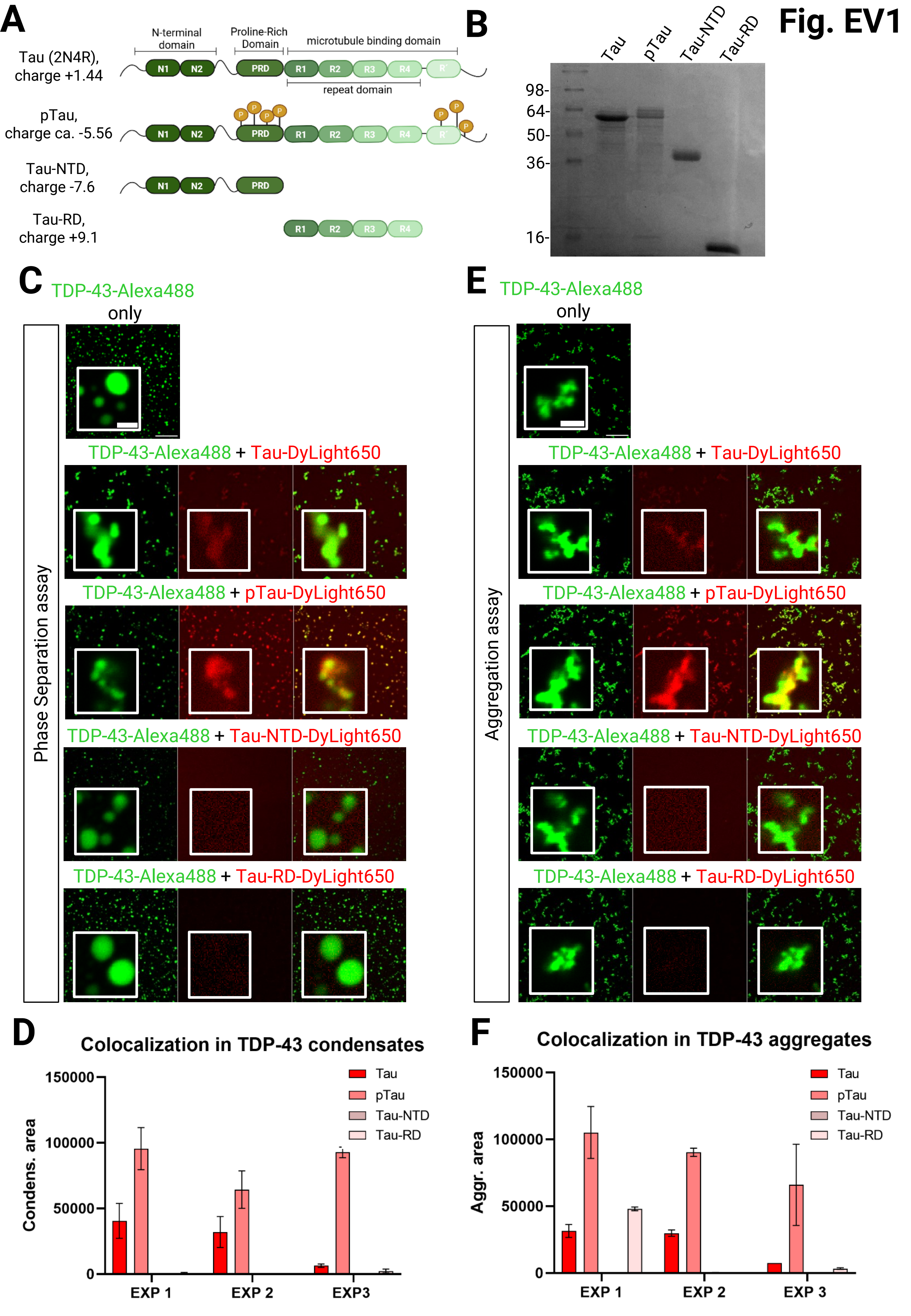
Fig. EV1
A
N-terminal domain
Proline-Rich Domain
microtubule binding domain
repeat domain
Tau (2N4R), charge +1.44
pTau, charge ca. -5.56
Tau-NTD, charge -7.6
Tau-RD, charge +9.1
N1
N2
PRD
R1
R2
R3
R4
R'
P
B
Tau
pTau
Tau-NTD
Tau-RD
98-
64-
50-
36-
16-
C
TDP-43-Alexa488 only
Phase Separation assay
TDP-43-Alexa488 + Tau-DyLight650
TDP-43-Alexa488 + pTau-DyLight650
TDP-43-Alexa488 + Tau-NTD-DyLight650
TDP-43-Alexa488 + Tau-RD-DyLight650
E
TDP-43-Alexa488 only
Aggregation assay
TDP-43-Alexa488 + Tau-DyLight650
TDP-43-Alexa488 + pTau-DyLight650
TDP-43-Alexa488 + Tau-NTD-DyLight650
TDP-43-Alexa488 + Tau-RD-DyLight650
D
Colocalization in TDP-43 condensates
Condens. area
150000
100000
50000
0
EXP 1
EXP 2
EXP3
Tau
pTau
Tau-NTD
Tau-RD
F
Colocalization in TDP-43 aggregates
Aggr. area
150000
100000
50000
0
EXP 1
EXP 2
EXP 3
Tau
pTau
Tau-NTD
Tau-RD

**Figure EV1: pTau, but not the Tau N-terminal domain or repeat domain, enriches in TDP-43 condensates and aggregates**

A. Scheme of recombinant Tau variants with their respective net charges; created with BioRender.com.

B. Coomassie-stained SDS-PAGE gel showing the molecular sizes of Tau, phosphorylated Tau (pTau), Tau N-terminal domain (Tau-NTD), and the Tau-RD fragment (R1–R4 repeat domains).

C. Confocal microscopy images of Alexa488-labeled TDP-43 (5 µM) in absence or presence of DyLight650-labeled Tau, pTau, Tau-NTD, or Tau-RD in a phase separation assay. Scale bar: 15 µm in overview and 3 µm in inset.

D. Colocalization of Tau, pTau, Tau-NTD, or Tau-RD within TDP-43 condensates quantified by measuring the area of condensates exhibiting overlapping green and far-red signals. Quantification was performed across (n = 3) biological replicates, and bar graphs show values ± SEM.

E. Confocal microscopy images of Alexa488-labeled TDP-43 (5 µM) in absence or presence of DyLight650-labeled Tau, pTau, Tau-NTD, or Tau-RD in an aggregation assay. Scale bar: 15 µm in overview and 3 µm in inset.

F. Colocalization of Tau, pTau, Tau-NTD, or Tau-RD within TDP-43 aggregates quantified by measuring the area of aggregates exhibiting overlapping green and far-red signals. Quantification was performed across (n = 3) biological replicates, and bar graphs show values ± SEM.

### Fig. EV2

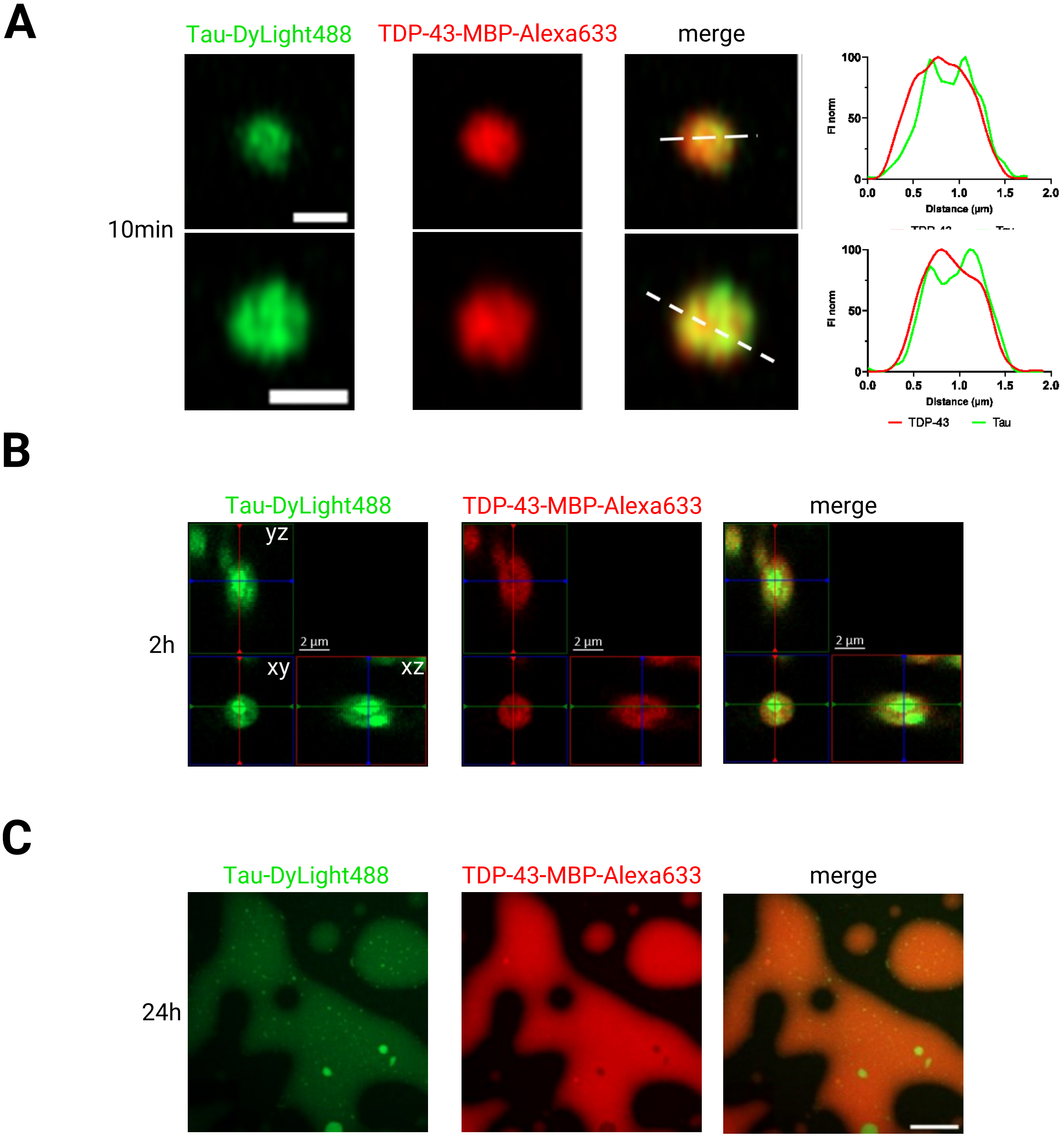


**Figure EV2: Early intra-droplet structures appear at the initial stages of multiphasic Tau/TDP-43 co-condensate formation**

A., B., and C. High resolution images using AiryScan of 1:1 DyLight488-labeled Tau with Alexa633-labeled TDP-43-MBP mixed condensates after 10 min (A), 2h (B) or (C) 24 h incubation. Scale bar: 1 µm (A), 2 µm (B) or 10 µm (C). In (A), dotted lines show the origin of line profiles indicated on the right, demonstrating demixing of Tau and TDP-43 within the same condensate at early timepoints.

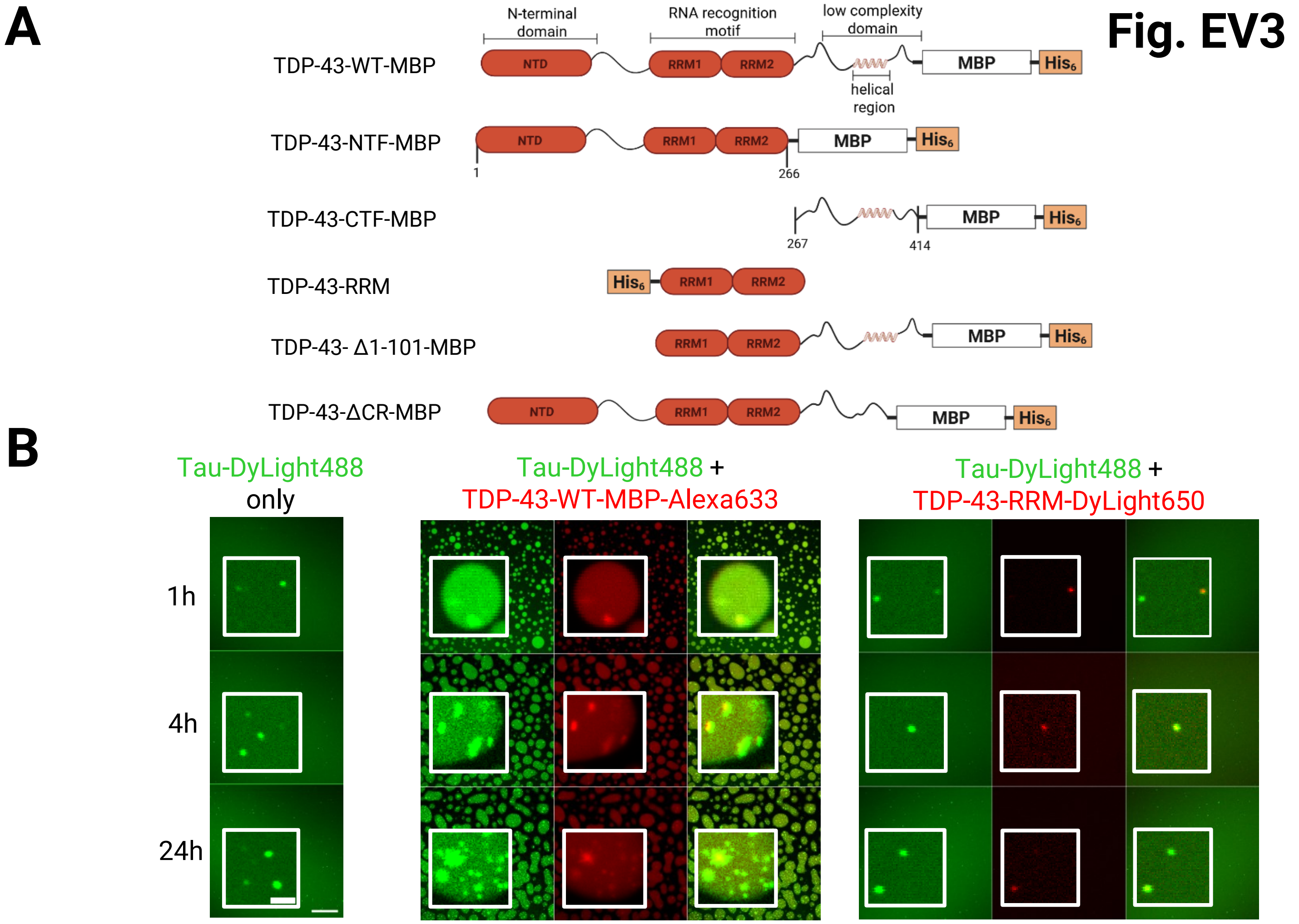


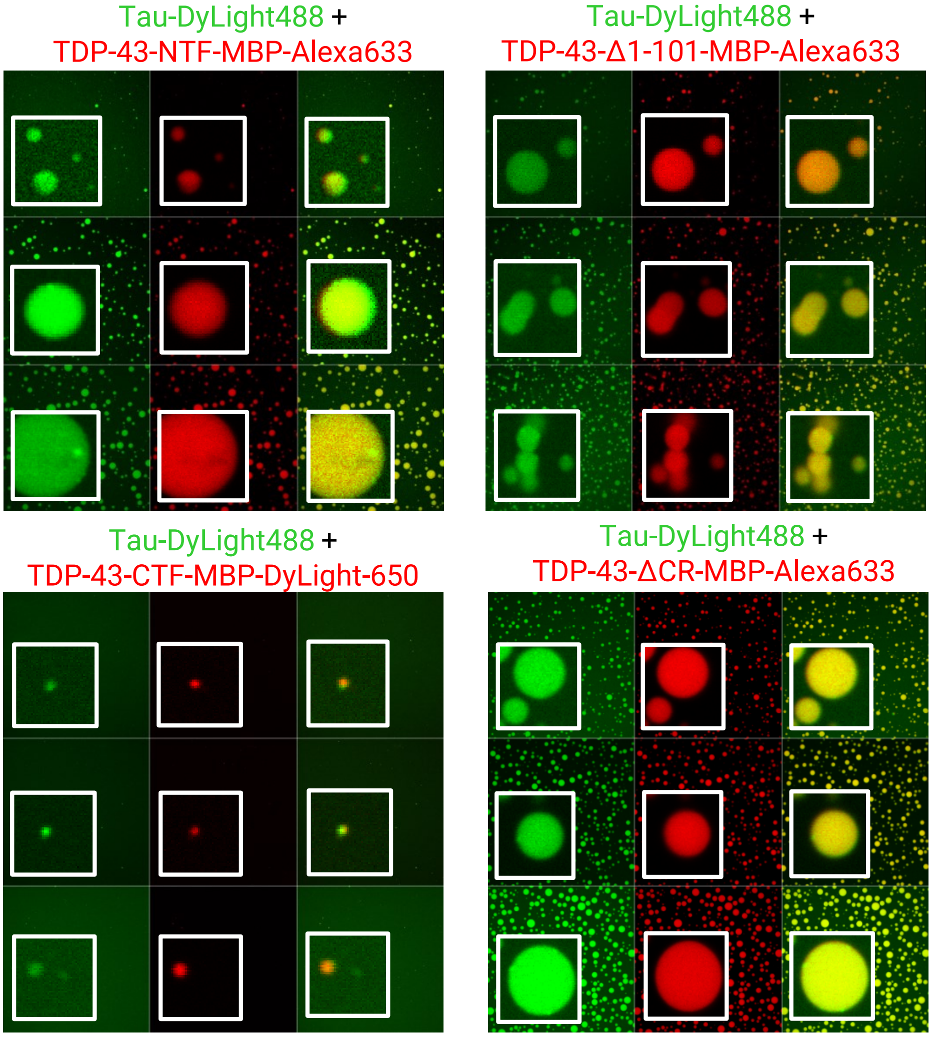


**Figure EV3: Distinct regions of TDP-43 contribute to Tau phase separation and the formation of multiphasic co-condensates**

A. Scheme of recombinant TDP-43 deletion mutants; created with BioRender.com.

B. Confocal microscopy images of 4 μM DyLight488-labeled Tau in equimolar presence of Alexa633-labeled TDP-43-WT-MBP, TDP-43-NTF-MBP, TDP-43-CTF-MBP, TDP-43-RRM, TDP-43-Δ1-101-MBP, or TDP-43-ΔCR-MBP at the indicated timepoints (1, 4, 24 h). Scale bar: 20 μm in overview and 2 μm in inset.

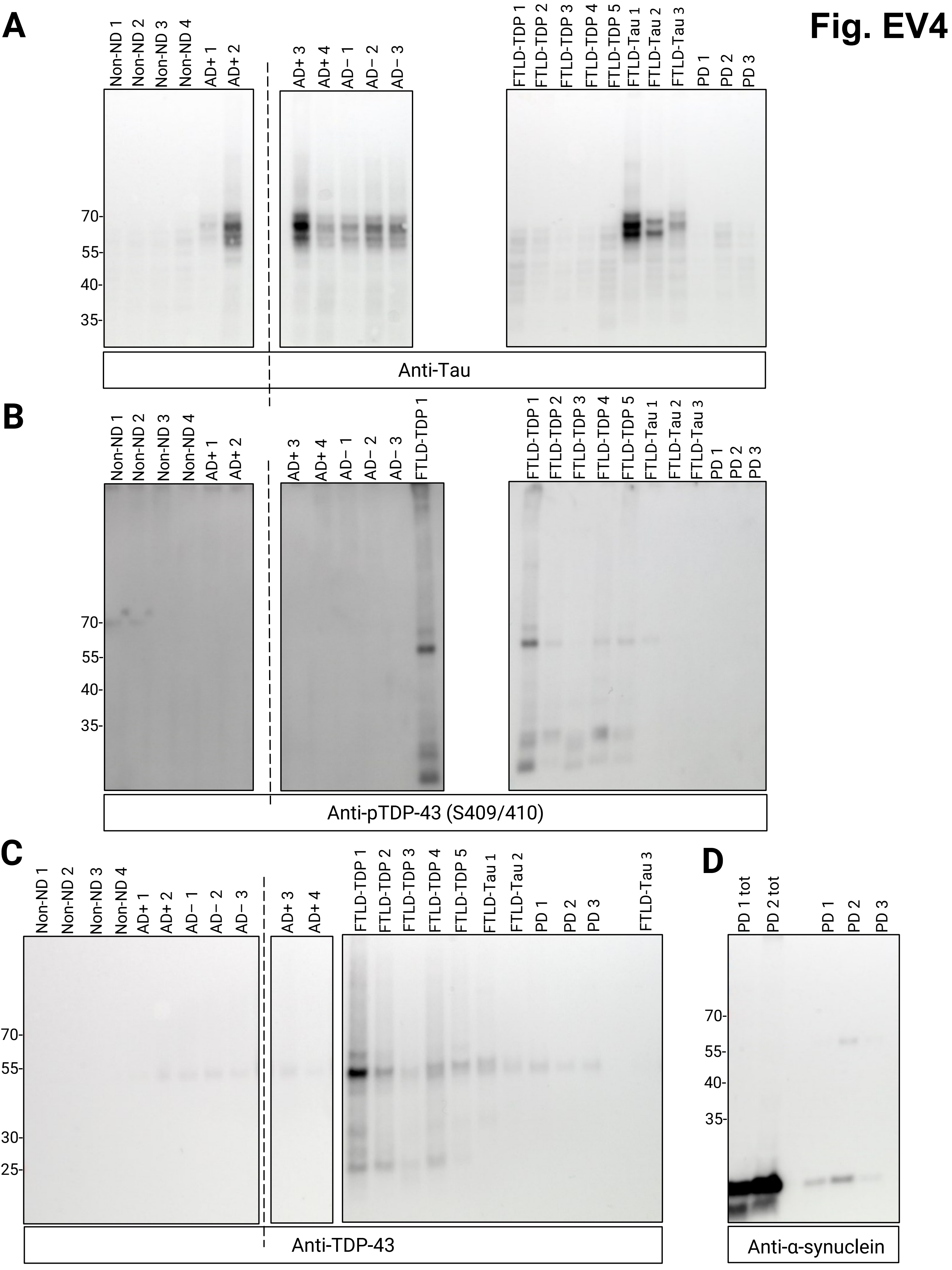


**Figure EV4: Western Blot characterization of SarkoSpin extracts from human patients**

Western blots of SarkoSpin fractions derived from the frontal cortex of Non-ND, AD+, AD–, FTLD-TDP, and FTLD-Tau patients, and from the cingulate cortex of PD patients, probed with the indicated antibodies. Dashed lines indicate divisions within the same blot.

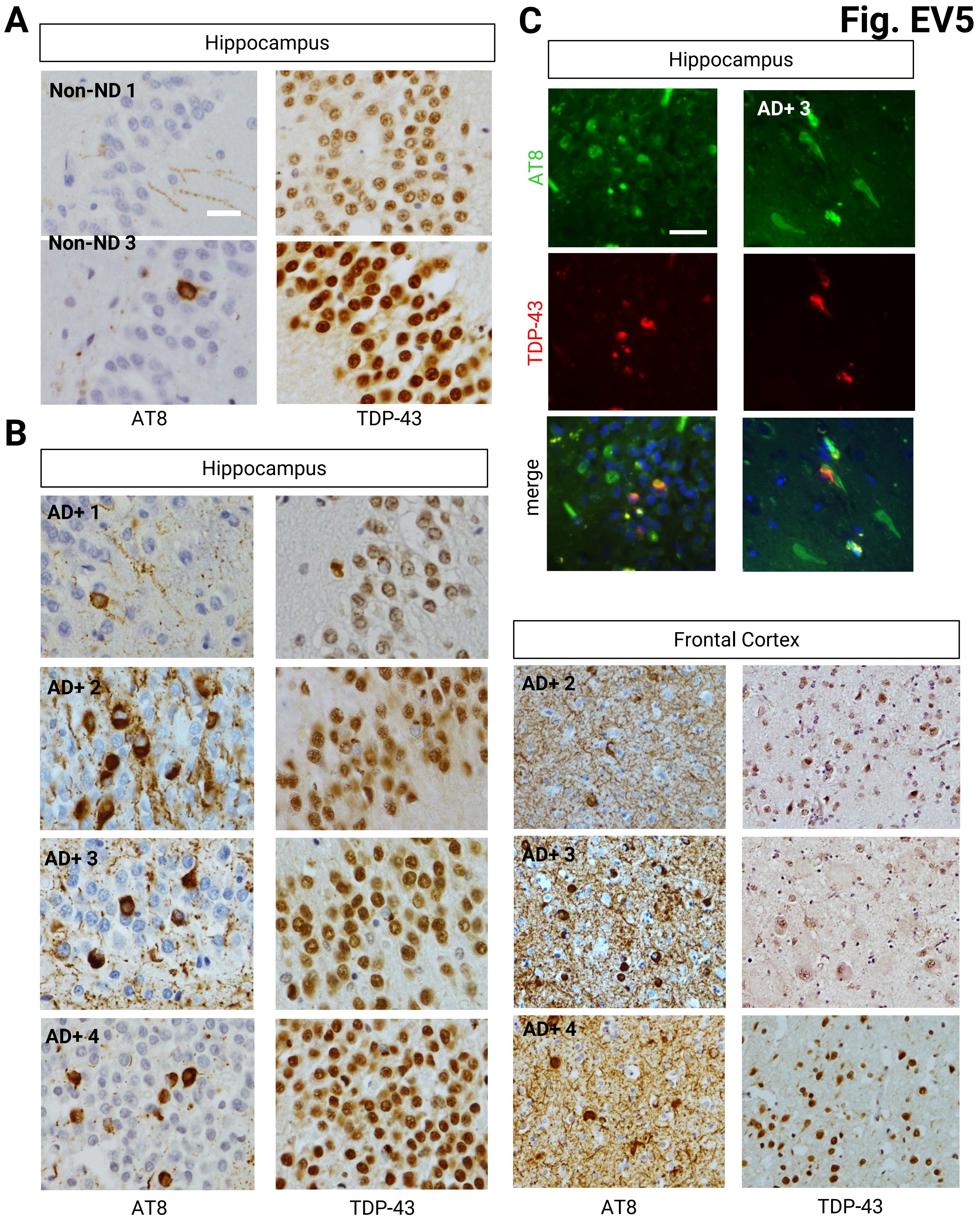


**Figure EV5: Characterization of patient-derived SarkoSpin extracts by immunohistochemistry and dual immunofluorescence**

A. and B. Immunohistochemical images of representative Non-ND and AD+ brain sections stained with antibodies against phosphorylated Tau (AT8) and TDP-43 in either the hippocampus or frontal cortex.

C. Double fluorescence immunostaining of two representative AD+ cases showing phosphorylated Tau (AT8, green) and TDP-43 (red) in the hippocampus.
