## Appendix for "Direct binding of TDP-43 and Tau drives their co-condensation, but suppresses Tau fibril formation and seeding"

#### Table of content

**Appendix Figure S1: Tau is more efficiently recruited into TDP-43 condensates and aggregates compared to α-synuclein**

**Appendix Figure S2: Crosslinks identification and Alpha Fold interaction prediction**

**Appendix Figure S3: Tau specifically promotes the aggregation of TDP-43 into high molecular weight (HMW) species**

**Appendix Figure S4: TDP-43, but not MBP or α-synuclein, promotes Tau condensation under low-salt, crowding-free conditions**

**Appendix Figure S5: Multiphasic condensate formation is independent of fluorescent labeling and crowding agent**

**Appendix Figure S6: Full-length TDP-43 suppresses Tau fibrillization at early time points**

**Appendix Figure S7: Characterization of patient-derived SarkoSpin extracts by immunohistochemistry**

**Appendix Figure S8: Representative microscopy images from Tau and TDP-43 seeding assays using patient-derived aggregates**

#### Appendix Fig. S1

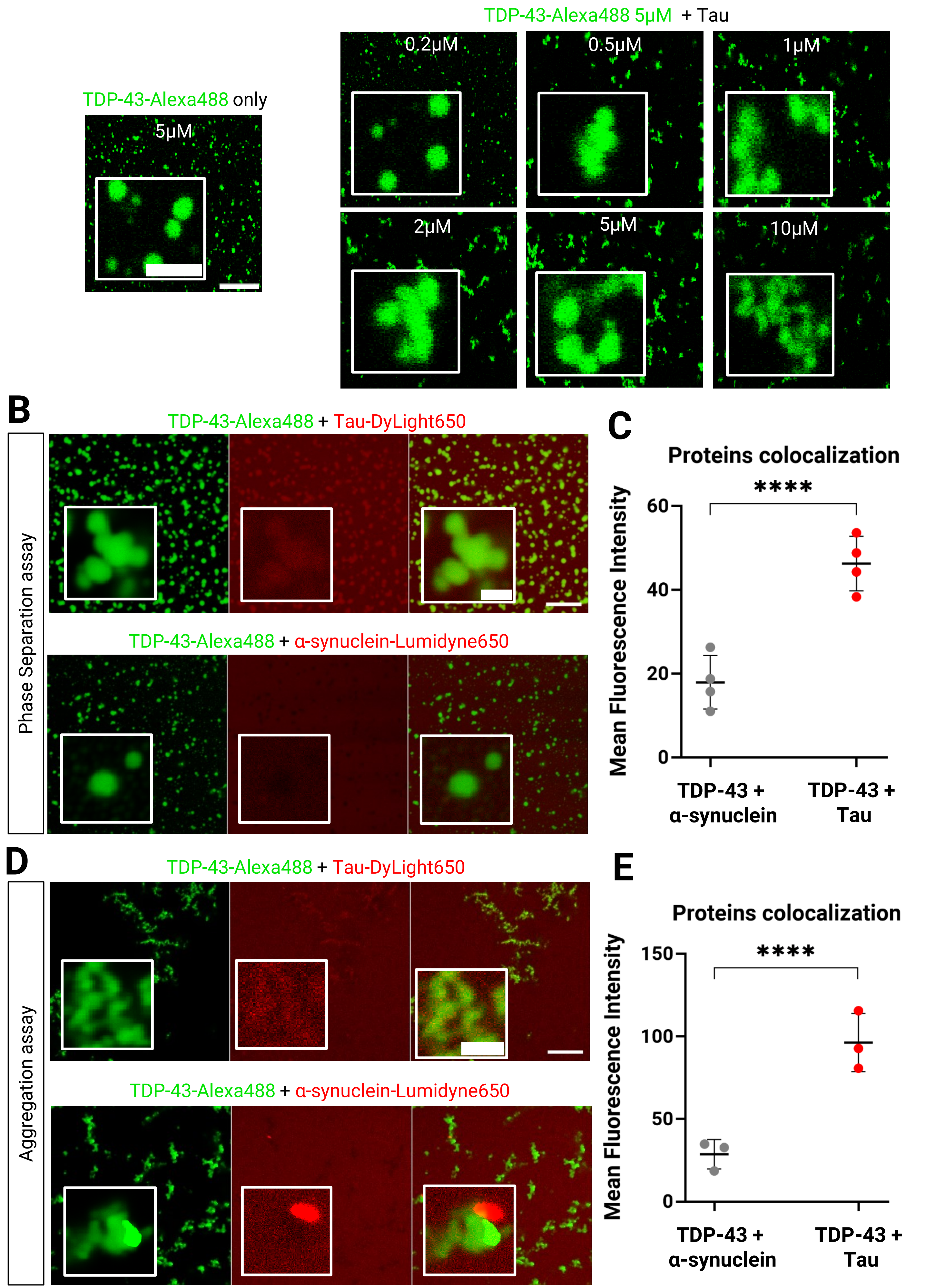

**Appendix Figure S1: Tau is more efficiently recruited into TDP-43 condensates and aggregates compared to α-synuclein**

A. Confocal microscopy images of Alexa488-labeled TDP-43 (5 µM) in presence of unlabeled Tau at the indicated concentrations (0.2, 0.5, 1, 2, 5, 10 µM) after ~30 min of TEV addition. Scale bar: 15 µm in overview and 5 µm in inset.

B. Condensates images of Alexa488-labeled TDP-43 (10 µM) with equimolar DyLight650-labeled Tau and Lumidyne650-labeled α-synuclein ~1h after TEV-cleavage. Scale bar: 20 µm in overview and 3 µm in inset. Linear adjustments to brightness and contrast were applied for visualization purposes.

C. Quantification of colocalization of DyLight650-labeled Tau and Lumidyne650-labeled α-synuclein within the Alexa488-labeled TDP-43 condensates in (n=4) biological replicates; values show the mean fluorescence intensity, and the bar graphs represents the mean ± SD; ****P < 0.0001 by unpaired t-test with Welch´s correction.

D. Confocal images of Alexa488-labeled TDP-43 aggregates (10 µM) formed in presence of DyLight650-labeled Tau or Lumidyne650-labeled α-synuclein (10 µM). Scale bar: 20 µm in overview and 3 µm in inset. Linear adjustments to brightness and contrast were applied for visualization purposes.

E. Quantification of the colocalization of DyLight650-labeled Tau and Lumidyne650-labeled α-synuclein within Alexa488-labeled TDP-43 aggregates in (n=3) biological replicates; values show the mean fluorescence intensity, and the bar graphs represents the mean ± SD; ****P < 0.0001 by unpaired t-test with Welch´s correction.

### Appendix Fig. S2

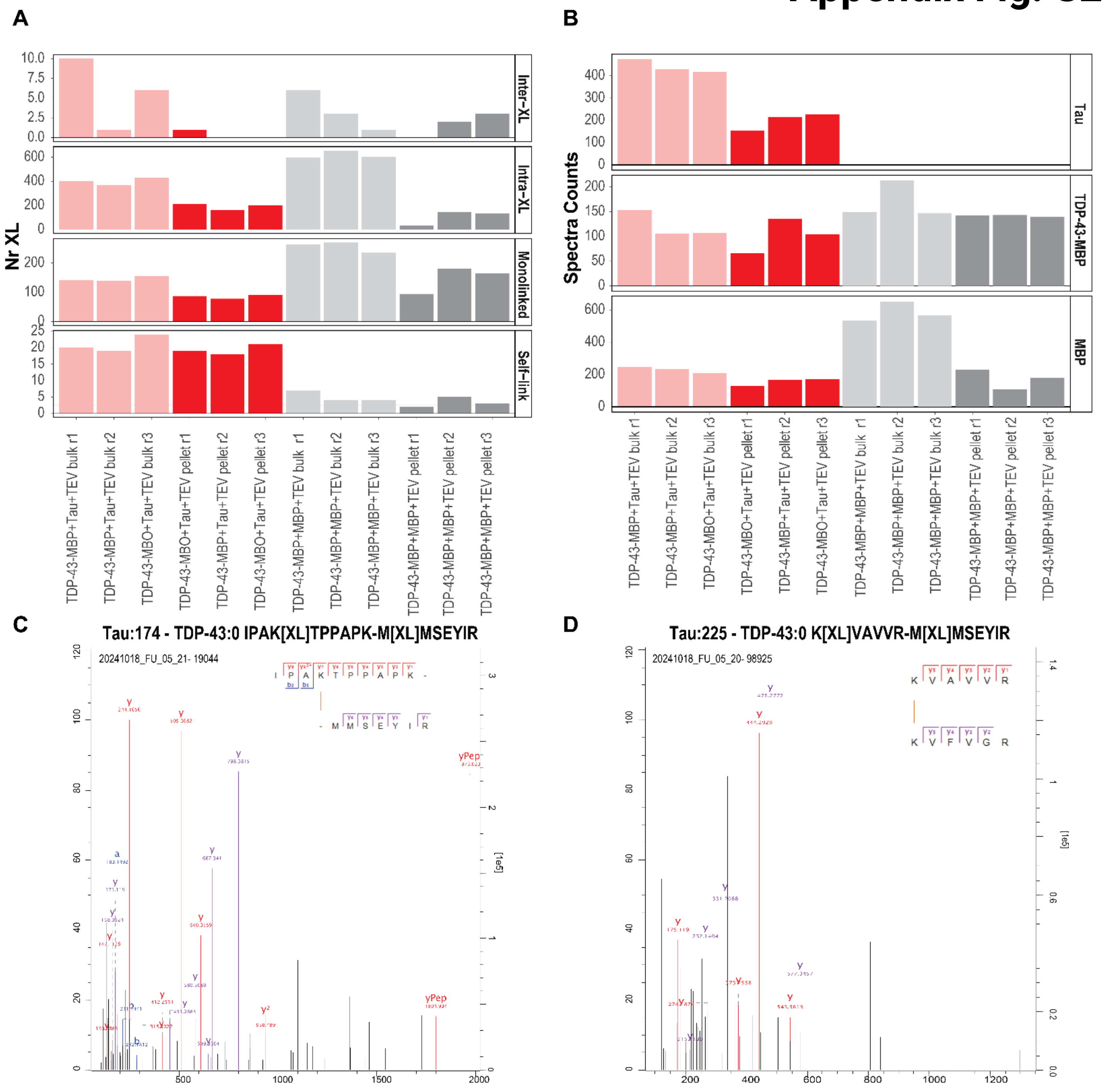


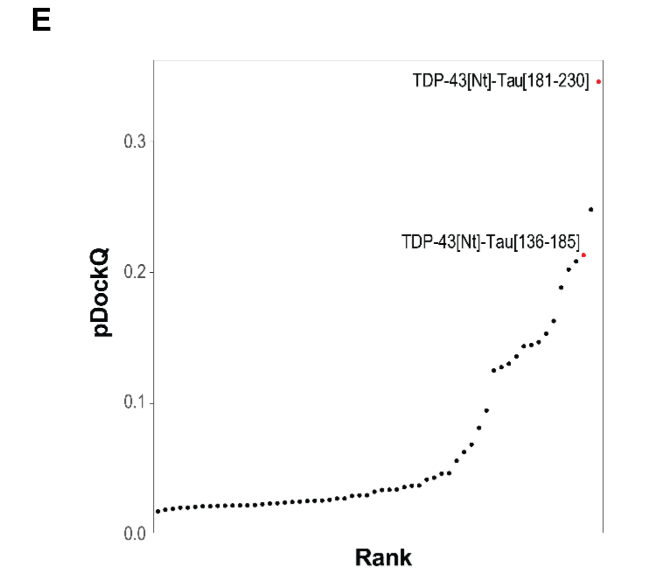


**Appendix Figure S2: Crosslinks identification and Alpha Fold interaction prediction**

A. Total number of spectra identified across all conditions containing at least one TDP-43 peptide crosslinked by DSS.

B. Total number of spectra counts for TDP-43, Tau and MBP across all measured conditions.

C. and D. MS spectra identifying the inter-protein crosslinked peptides TDP-43 0 – Tau 174 and TDP-43 0 – TAU 225, respectively.

E. Ranked AlphaFold models based on pDockQ scores for all 60 pairwise permutations of TDP-43 and TAU fragments. Models containing the crosslinked peptide pairs TDP-43 0 – Tau 174 and TDP-43 0 – Tau 225 are highlighted.

### Appendix Fig. S3

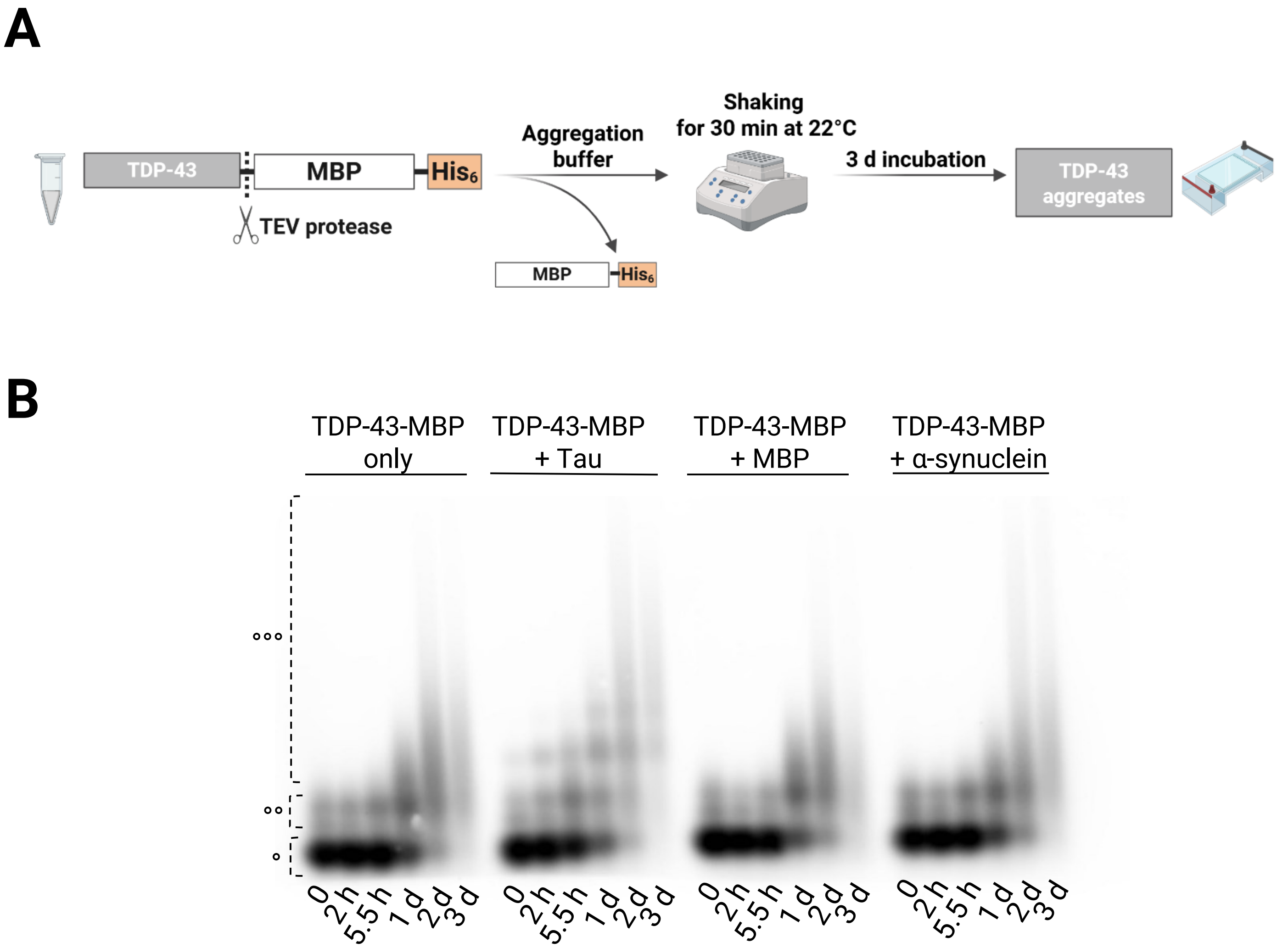


**Appendix Figure S3: Tau specifically promotes the aggregation of TDP-43 into high molecular weight (HMW) species**

A. Scheme of TDP-43 aggregation assay with TEV cleavage for SDD-AGE experiment; created with BioRender.com.

B. SDD-AGE of TDP-43 (2 μM) in the presence of the indicated protein (2 μM) after TEV cleavage, agitation and incubation for the presented time period (h = hours, d = days); TDP-43 was visualized by immunoblotting and the middle vertical black line divides two blots which derive from the same experiment and were processed in parallel. °°° = high molecular weight (HMW) species, °° = oligomers and ° = monomers.

### Appendix Fig. S4

A

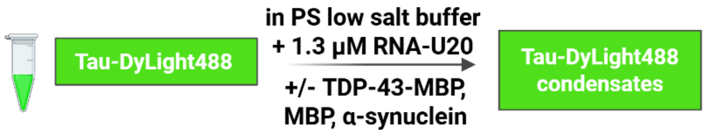


B

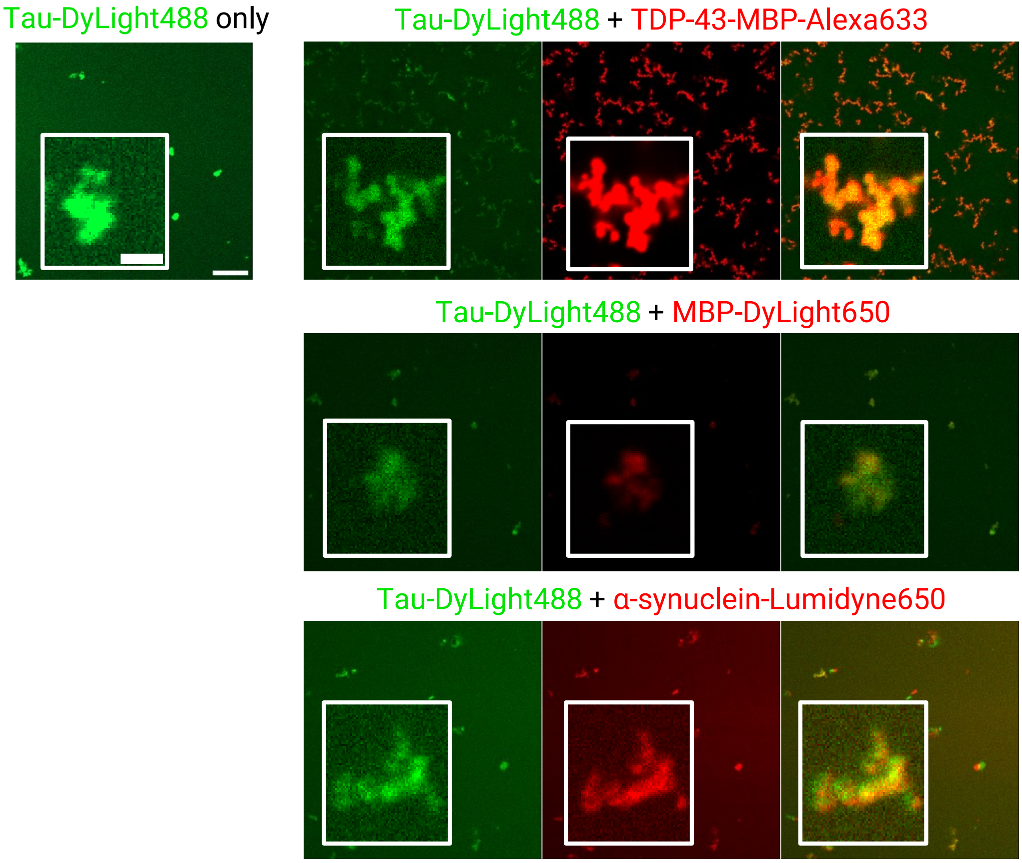


**Appendix Figure S4: TDP-43, but not MBP or α-synuclein, promotes Tau condensation under low-salt, crowding-free conditions**

A. Schematic representation of Tau phase separation assay in the presence of RNA; created with BioRender.com.

B. Confocal images of DyLight488-labeled Tau (5 µM) only, or in the presence of Alexa633-labeled TDP-43-MBP, DyLight650-labeled MBP or Lumidyne650-labeled α-synuclein at equimolar concentration after ~1 h of RNA-U20 addition. Scale bars: 15 µm in overview and 2 µm in inset.

### Appendix Fig. S5

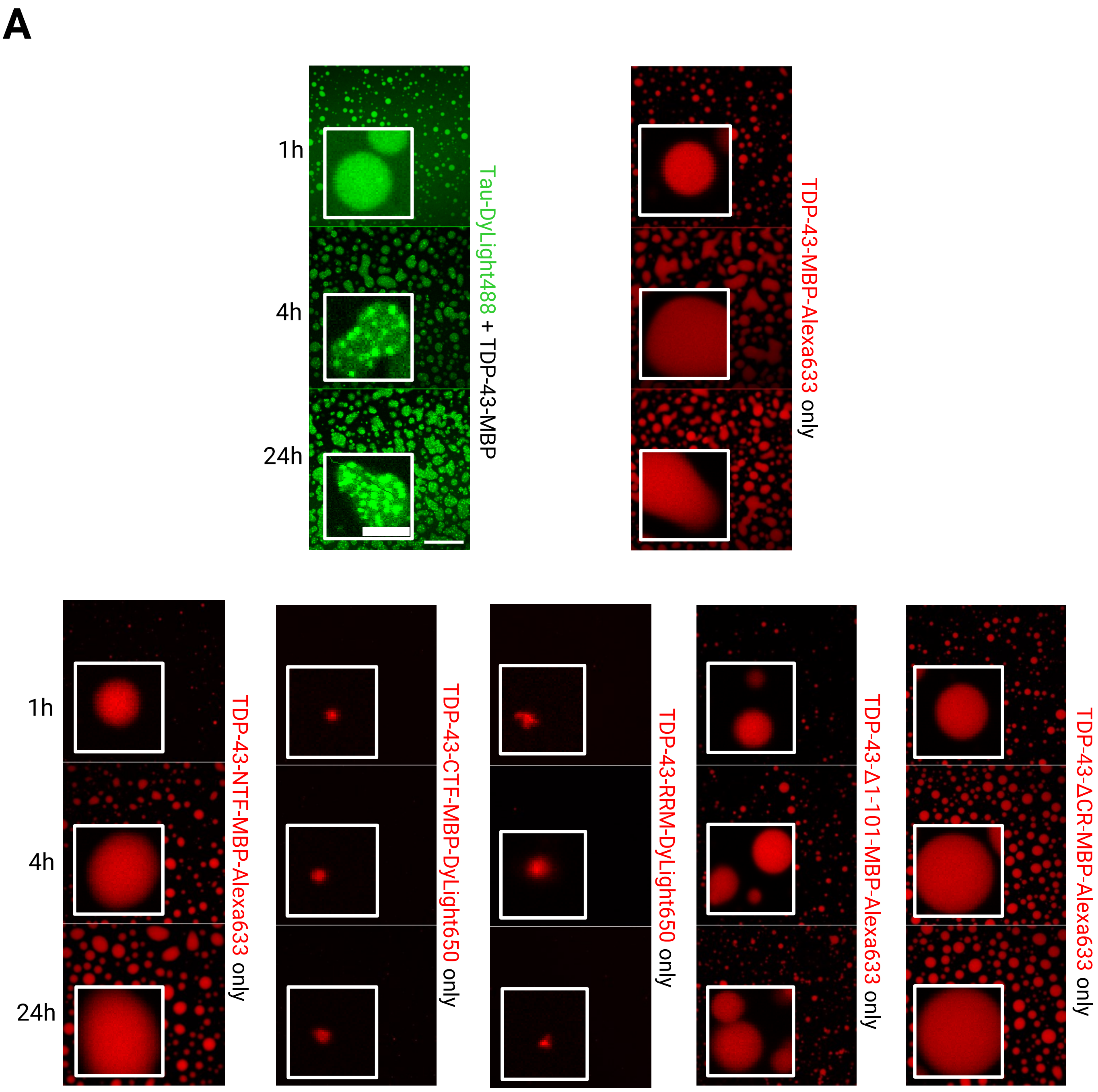


**Appendix Figure S5: Multiphasic condensate formation is independent of fluorescent labeling and crowding agent**
A. Confocal images of unlabeled TDP-43-MBP, Alexa633-labeled TDP-43-WT-MBP, TDP-43-NTF-MBP, TDP-43-CTF-MBP, TDP-43-RRM, TDP-43-Δ1-101-MBP, or TDP-43-ΔCR-MBP in the presence of 10 % PEG. Scale bars: 25 µm in overview and 4 µm in inset.

### Appendix Fig. S6

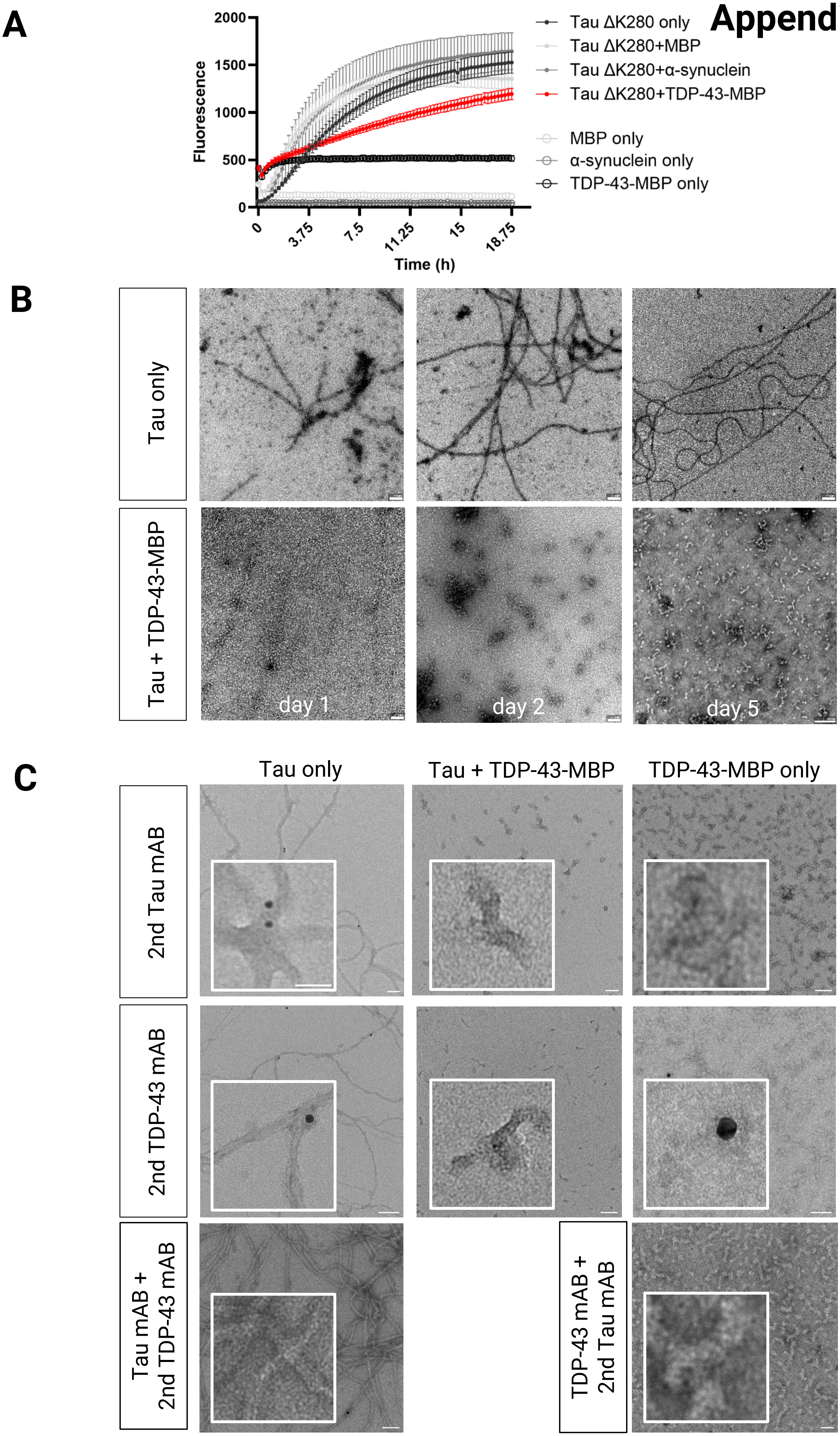

**Appendix Figure S6: Full-length TDP-43 suppresses Tau fibrillization at early time points**

A. Graph showing raw data ThT fluorescence measurements over 18 h.

B. TEM images of Tau or Tau + TDP-43-MBP (both 50 µM) at the indicated time points (day 1, 2, 5). Scale bar: 0.2 µm.

C. Representative TEM images of Tau-only, Tau + TDP-43, and TDP-43-only conditions stained using control secondary antibodies: secondary Tau antibody alone (2nd Tau mAB), secondary TDP-43 antibody alone (2nd TDP-43 mAB), or in combination with primary TDP-43 or Tau antibodies, respectively (TDP-43 mAB + 2nd Tau mAB; Tau mAB + 2nd TDP-43 mAB). Scale bars: Scale bar: 0.05 µm and 0.03 µm in inset.

### Appendix Fig. S7

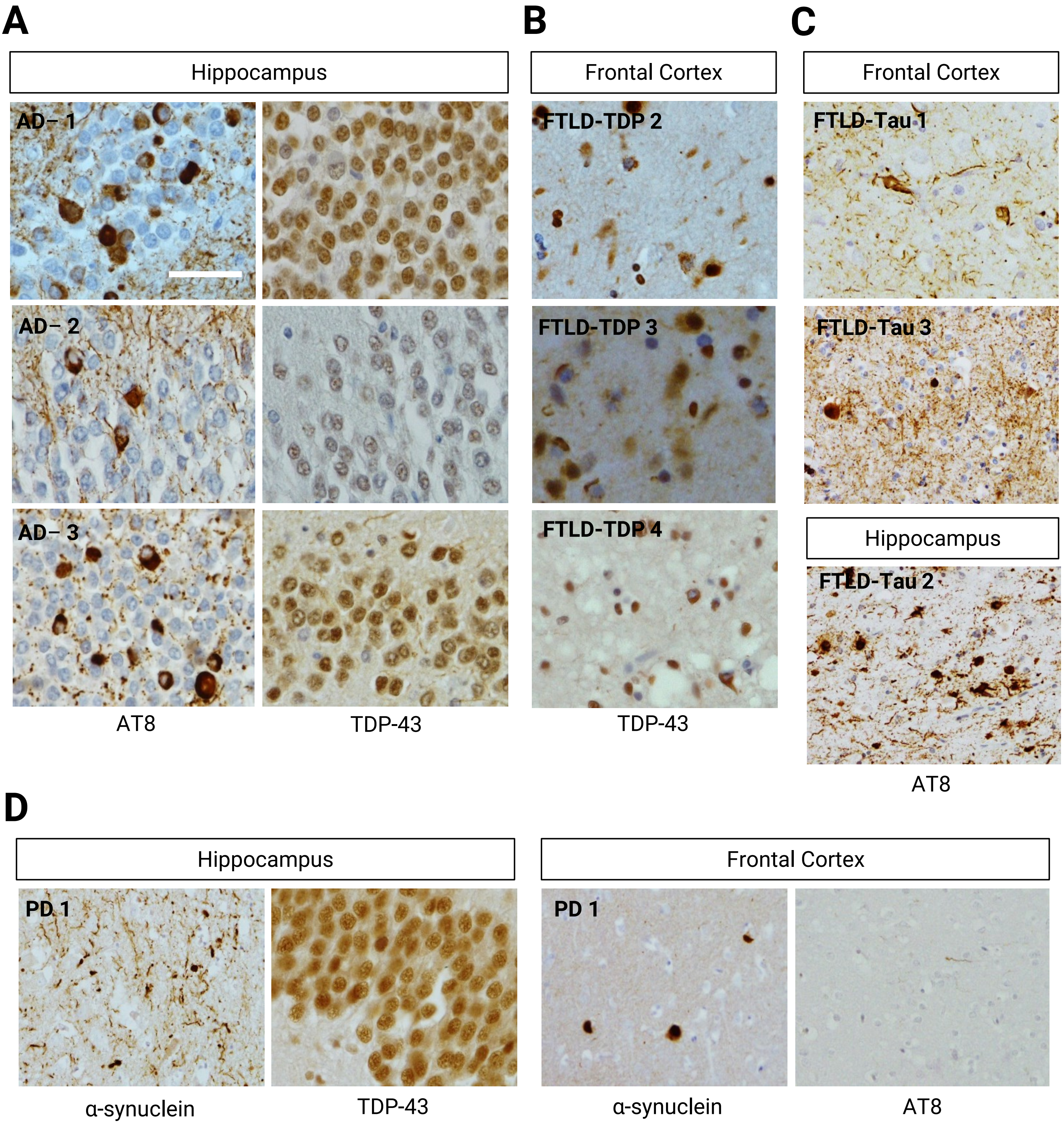


**Appendix Figure S7: Characterization of patient-derived SarkoSpin extracts by immunohistochemistry**

A. Immunohistochemical images of representative AD−, FTLD-TDP, FTLD-Tau and PD brain sections stained with antibodies against phosphorylated Tau (AT8), TDP-43 and α-synuclein in either the hippocampus or frontal cortex.

Appendix Fig. S8

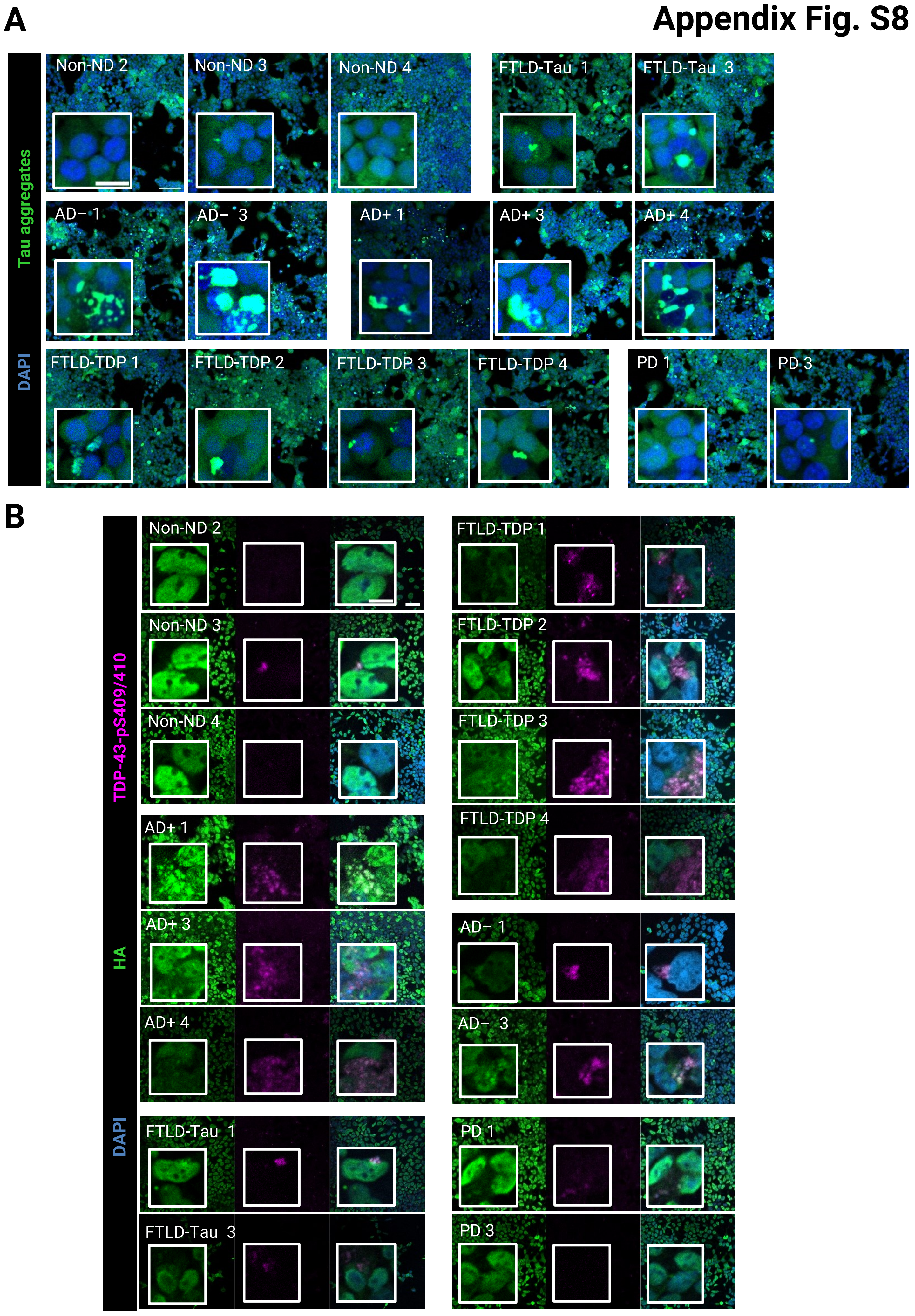

**Appendix Figure S8: Representative microscopy images from Tau and TDP-43 seeding assays using patient-derived aggregates**

A. Representative confocal images of cytosolic Tau aggregates (green) formed after seeding the cells with Non-ND 2, Non-ND 3, Non-ND 4, FTLD-Tau 1, FTLD-Tau 3, AD− 1, AD− 3, AD+ 1, AD+ 3, AD+ 4, FTLD-TDP 1, FTLD-TDP 2, FTLD-TDP 3, FTLD-TDP 4, or PD 1 and PD 3 SarkoSpin extracts. Scale bar: 100 µm in overview and 20 µm in inset.

B. Representative confocal images of cytosolic pS409/410-positive TDP-43 neoaggregates in HEK293 cells, after transfection with Non-ND 1, FTLD-TDP 5, AD+ 2, AD− 2, FTLD-Tau 2, or PD 2 SarkoSpin patient extracts. DAPI is depicted in blue, HA staining in green, and pS409/410 in magenta. Scale bar: 50 µm in overview and 10 µm in inset.
